# Proposal for a Global Adherence Scale for Acute Conditions (GASAC): a prospective cohort study in two Emergency Departments

**DOI:** 10.1101/598409

**Authors:** Mélanie Sustersic, Aurélie Gauchet, Amélie Duvert, Laure Gonnet, Alison Foote, Céline Vermorel, Benoit Allenet, Jean-Luc Bosson

**Author notes:** Corresponding author: (MS). These authors contributed equally to this work (Amélie Duvert, Laure Gonnet).

## Abstract

**Background:** Adherence in the context of patients with acute conditions is a major public health issue. It is neglected by the research community and no clinically validated generic scale exists to measure it.

**Objective:** To construct and validate a Global Adherence Scale usable in the context of Acute Conditions (GASAC) that takes into account adherence both to advice and to all types of prescriptions that the doctor may give. To measure adherence and to study its determinants.

**Materials and method:** We based the construction of the GASAC questionnaire on a theoretical model and a literature search. Then, between 2013 and 2014, we validated it in a prospective observational study in two hospital emergency departments. Patients were contacted by phone about one week after their consultation to answer several questionnaires, including GASAC and the Girerd self-administered questionnaire about medication adherence as a control.

**Results:** GASAC consists of four adherence subscales: drug prescriptions; blood test/ radiography prescriptions; lifestyle advice and follow-up instructions. An analysis of the 154 sets of answers from patients showed that the GASAC drug subscale had satisfactory internal coherence (Cronbach’s alpha = 0.78) and was correlated with the Girerd score, as was GASAC as a whole (p<0.01)). The median score was 0.93 IQR [0.78-1] for a maximum value of 1 (n = 154). In multivariate analysis, infection was more conducive of good adherence (cut off at ≥ 0.8; n=115/154; 74.7% [67.0-81.3]) than trauma (OR 3.69; CI [1.60-8.52]). The Doctor-Patient Communication score (OR 1.06 by score point, CI [1.02-1.10]) also influenced adherence.

**Conclusions:** GASAC is a generic score to measure all dimensions of adherence in emergency departments for clinical research and the evaluation of clinical practice. The level of adherence was high for acute conditions and could be further improved by good Doctor-Patient Communication.

## Introduction

Adherence is an important component in the assessment of the quality of healthcare. A review of the literature on the evaluation of adherence found 90 articles published in the last 10 years [1]. For the World Health Organization (WHO) “Optimizing medication adherence would have more impact in terms of global health than the development of new drugs [2].” Non- adherence leads to a considerable waste of health resources (e.g. unnecessary hospitalizations, unused drugs) and is a cause of avoidable morbidity and mortality [3,4].

While chronic diseases (hypertension, asthma, HIV etc.) have received much attention [2,5,6], adherence following an acute condition (AC) has been relatively little studied. Most of the studies in this field have been performed for a particular chronic disease and many authors mainly studied adherence to drug prescriptions [7–13]. In only a few cases did they look at adherence in a broader sense [4,14–16]. The most widely used scale, considered a reference in the field [17,18], the Morisky score, measures drug adherence for chronic conditions [19,20]. It is sometimes criticized for its lack of discrimination [21]. The Girerd adherence scale, although initially created for chronic disease, has several advantages: the items that constitute it are compatible with its use in acute conditions and there is a validated French version [10,11].

AC are the most frequent reason for consultations in both general practice [22] and Emergency Departments (ED). Primary care services are overcrowded and the management of AC is becoming a major public health issue [23]. The period after a consultation for an AC is one of high vulnerability for patients [24]. They run the risk of further health deterioration, may suffer from a misdiagnosis [23] or even experience side effects from a newly prescribed drug [24]. Immediate vital (e.g. some infections) and/or long-term outcomes (eg. sequels of an ankles sprain) might possibly arise and could potentially affect the prognosis [23]. In such a context, adherence to the doctor’s advice and to all types of prescriptions given is key to the success of any treatment [25]. Any aspects of the patient’s behaviour (acting upon prescriptions for X-rays and laboratory tests, respecting appointments with specialists etc.) could be as important as adherence to medications, which is already spontaneously high [3,26,27].

The notion of adherence uniquely to prescriptions for medication is insufficient [27,28]. Although some authors defined adherence “as the extent to which a patient’s behaviour (in terms of taking medication, following a diet, modifying habits, or attending clinics) coincides with medical or health advice” [4], there is still no method to measure adherence that includes all aspects of a patient’s behaviour after a consultation. There lacks a standardized tool that is well adapted to clinical research [29] and assessment of health-care quality [1,17], and none suitable for the context of AC [30,31]. The question of how best to measure adherence is still open [2].

A generic scale would be useful to analyse the relationship of between adherence to and other outcomes such as Doctor-Patient Communication (DPC) or satisfaction, and to quantify and compare the impact of measures introduced to improve adherence, such as Patient information leaflets [30,31].

Our objective was to create a Global Adherence Scale usable in the context of an Acute Condition (GASAC) based on a theoretical model describing the various dimensions of patient behaviour following a consultation [15] and the results of a literature search. Then, to validate it in two hospital emergency departments and analyse its determinants.

## Materials and methods

### Literature search

We searched the Medline database using the following Mesh terms: patient compliance, adherence AND scale, tool, assessment, measures or questionnaires, in various combinations. We also consulted the Embase and PsycInfo databases, and the Cochrane library in English.

Our search filter covered the period from 1985 to 2014. Only meta-analyses, randomized controlled trials, and reviews of the literature were retained. In addition we searched English (NHS) and US (Agency for Healthcare Research and Quality, AHRQ) institutional databases on quality of care assessment and books on the field. Two doctors independently screened titles and if necessary abstracts for all types of articles pertinent to adherence in the context of acute conditions. A manual search was also conducted from the bibliographies of promising articles. Since our literature search did not find specific articles for AC, we based the elaboration of our scale on: 1/ a previously constructed theoretical model, itself based on the literature, 2/ commonly used definitions of adherence [2,4] and 3/ on an analysis of the literature so as not to overlook any dimension in our scale [32, 36].

## Questionnaire development

### Theoretical model used

Analysis of adherence behaviour from a strictly medical point of view is insufficient [32]. For this reason, we based the construction of our scale on an existing model recently developed by a multidisciplinary team [15] using a multifactorial approach, as recommended by studies in psychology and sociology [5,32]. It also helped us to avoid the pitfalls of vague terminology [31], poor construction of the scoring system and redundancy between outcomes [15]. This model describes the four aspects of a patient’s behaviour following a consultation: taking medications as instructed, following prescriptions for evaluations and tests (radiography, blood tests, appointments with specialists), making appropriate lifestyle changes (i.e. diet, stopping smoking, physical activity, alcohol consumption) and when to engage the healthcare system for worsening or reoccurring symptoms and follow-up. Respecting the model’s categories, we constructed a rigorous scale with pertinent items. Moreover the model assisted us in studying the determinants of adherence such as DPC and satisfaction, also defined in the model, with a solid theoretical foundation.

### Requirements of the new scale

The scale needed to respect the following criteria: usable in routine practice, independent of any particular clinical situation, self-reported by the patient, easy to understand, easily evaluable, brief, respectful of the patient’s privacy, possible to be completed by or together with carers where necessary, validated and reliable [20]. Self-assessment by the patient was considered the method of choice as it is fast, inexpensive, non-invasive and can potentially help detect the underlying reasons for non-adherence [21,37]. In practice certain authors consider it to be the most suitable method for assessing health improvement [20] although it might sometimes suffer from self-deception or dishonest answers. An objective measure such as pill counts, sometimes used in assessing drug adherence, was not feasible in the broad context of assessing other types of behaviour.

## Validation of the questionnaire

### The pilot study

The first version was tested in a pilot study on 30 patients whatever the pathology diagnosed. Immediately after the consultation the patient was given the questionnaire to complete, followed by an additional page about their understanding of the questionnaire and open remarks.

### Sample size calculation

We based our calculation on the general adherence literature according to a systematic Cochrane Library review [27]. To measure the impact of any intervention on adherence we needed to obtain a minimum of 75 completed questionnaires for each group. As we intended to include patients with either a non-severe trauma i.e. ankle sprain, or a medical indication i.e. an infection, this meant the minimum sample size for our study was 150.

Assuming good adherence (estimated at 85%) [3] a sample size of 150 patients would provide an accuracy of measurement, with a 95% confidence interval (CI) of ±6%. The inclusion of 150 patients would also allow a calculation of the Cronbach coefficient with a satisfactory level of precision (Cronbach alpha with one-sided 95%; CI = 0.75 for three items of the pharmacological subscale and a coefficient estimated at 0.8).

Allowing for 20% patients potentially being lost to follow-up we required 180 patients in total. We stopped inclusions when this number was reached.

### Design

A two-centre prospective observational study was conducted from November 2013 to May 2014 in the emergency departments of two hospitals. The study was approved by the regional ethics committee (IRB n° 5891 on 31-Oct-2013).

Physicians who regularly worked in the ED of the two establishments were contacted and voluntarily participated. The physician briefly presented the study (orally and in a patient information letter) and proposed participation to all consecutive adults and children (>15 and accompanied by an adult) diagnosed with a common traumatic or infectious acute condition (ankle sprain or infectious colitis, pyelonephritis, diverticulitis, prostatitis or pneumonia). These acute conditions were chosen from those most commonly seen in primary care [22] and which usually require medication, prescriptions for specialist evaluation or tests, and/or advice and follow-up instructions. We excluded patients whose care led to a hospital stay of more than 48 hours.

The patient information letter explained the broad aims of the study: to help develop tools to measure the quality of care. Details were not given so as to reduce any self-selection bias. If the patient agreed to participate, they signed a written informed consent. If they declined to participate, this was recorded. Physicians included patients in the study by completing a short inclusion-case report form, describing the patient’s baseline and socio-demographic characteristics.

Patients were contacted by telephone between 7 and 10 days after the consultation by a study investigator who did not participate in patient recruitment. They were asked the series of questions from the Girerd scale (yes/no) and then the GASAC questions (scored on a Likert scale of 1 to 4). Next they were asked a series of questions about DPC, about patient satisfaction and some additional questions about the intentional or non-intentional nature of their behaviour (not counted in the score).

### Statistical analysis

Statistical analysis was performed using Stata Version 13.0 (Stata Corp, College Station, Texas). Statistical tests were performed with a significance level of 0.05. Qualitative variables are described by frequency and percentages, and quantitative variables using medians and IQR [25th and 75th percentiles]. The internal consistency of the GASAC score items was assessed by Cronbach’s alpha [34]. For quantitative variables, we used the Mann-Whitney test to compare two groups, or the Kruskal-Wallis test to compare more than two groups (non-parametric tests). For qualitative variables, we used the Chi-squared test. Finally, we conducted a multivariate logistic regression to identify factors associated with a “high” adherence according to the distribution of the histogram. To study the determinants of adherence we had to dichotomise the variables, so patients with a GASAC score ≥ 0.8 were classed as “highly-adherent”, whereas those with a score < 0.8 were classed as “poorly-adherent”. All patient characteristics, the level of information, satisfaction and DPC score with p<0.2 in univariate analysis were included into the full model. The final model was obtained by a manual step-wise logistic regression. The correlation between GASAC and the DPC, and correlation between subscales of GASAC scores was explored by calculating Spearman’s rho.

## Results

### Literature search

Our search extracted 845 records, including 80 reviews. Among these, neither of the two doctors found any reviews or original articles that dealt with an acute condition, nor with global adherence. Concordance was 100%. Among the reviews, four dealt with adherence to exercises (e.g. for musculoskeletal disease); one with showing up at a mammography appointment and one with keeping to a diet. The rest concerned drug adherence in chronic diseases (HIV, psychiatric disease, diabetes). Due to the lack of specific articles we consulted those dealing with adherence from a general point of view, non-specific to any given disease, and searched their bibliographies manually. We used scales measuring the quality of care from articles, the institutional websites and books [32–36]. Thus while the literature did not provide us with any suitable scales it helped us to be more precise and to better formulate our questions. We profited from the accumulated knowledge of the multidisciplinary research team that included experts in “adherence” (AG, BA).

### Description of the new scale

GASAC incorporates four adherence subscales: drug prescriptions (3 items); laboratory tests and/or radiography prescriptions (one item); lifestyle advice (one item) and instructions on when, or if, to consult a medical professional again (one item). These 6 items were each rated from 1 to 4 (1 = no, 2 = rather not, 3 = rather yes, 4 = yes). As not all questions were relevant to all patients (not all patients received prescriptions and /or instructions) the final score was expressed as the ratio, between 0 and 1, of the patient’s answers to the questions actually posed by the study investigator and the maximum possible score from this number of questions (Table 1). The mean time to complete the questionnaire was 3 minutes.

**Table 1.**
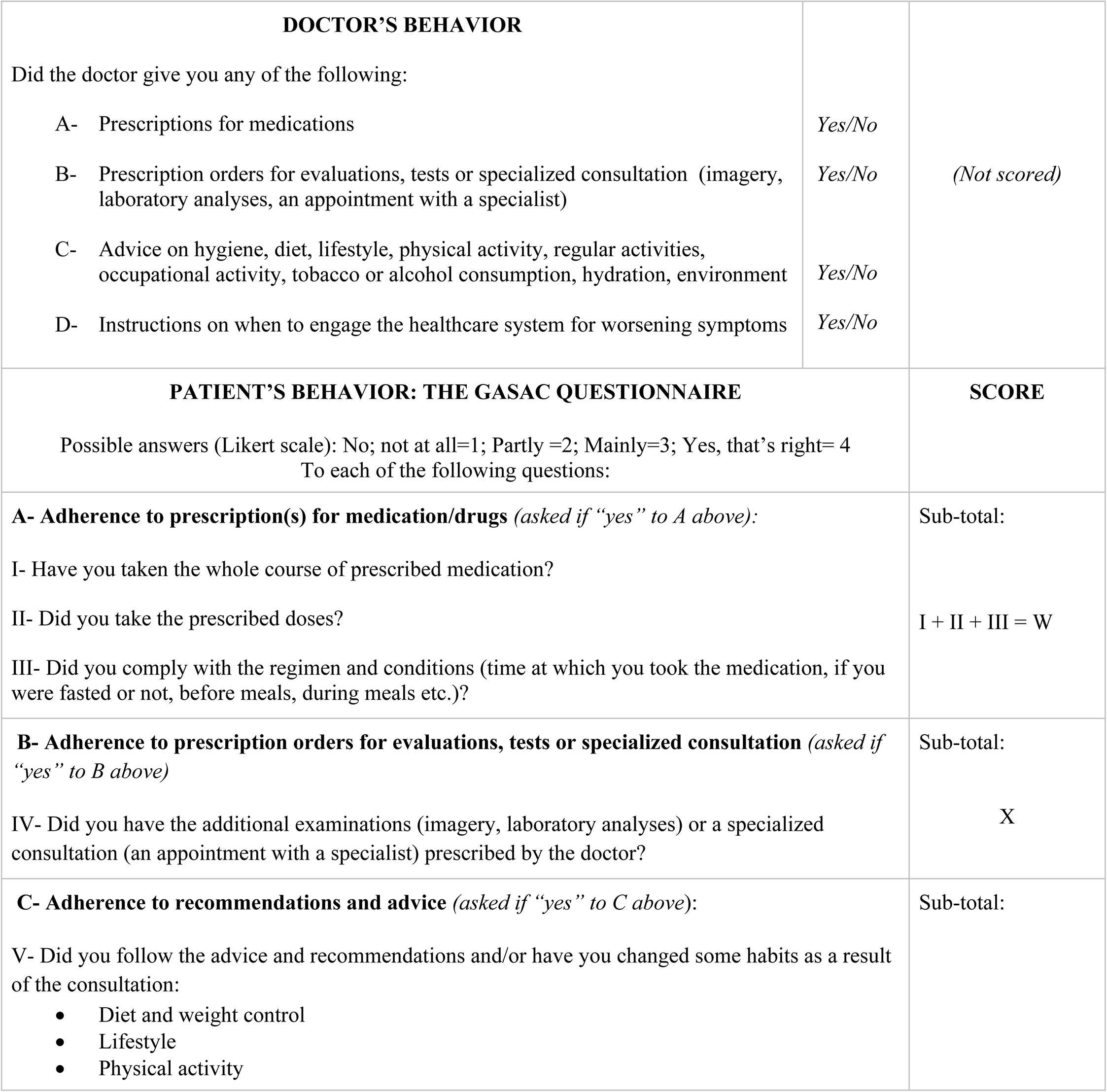
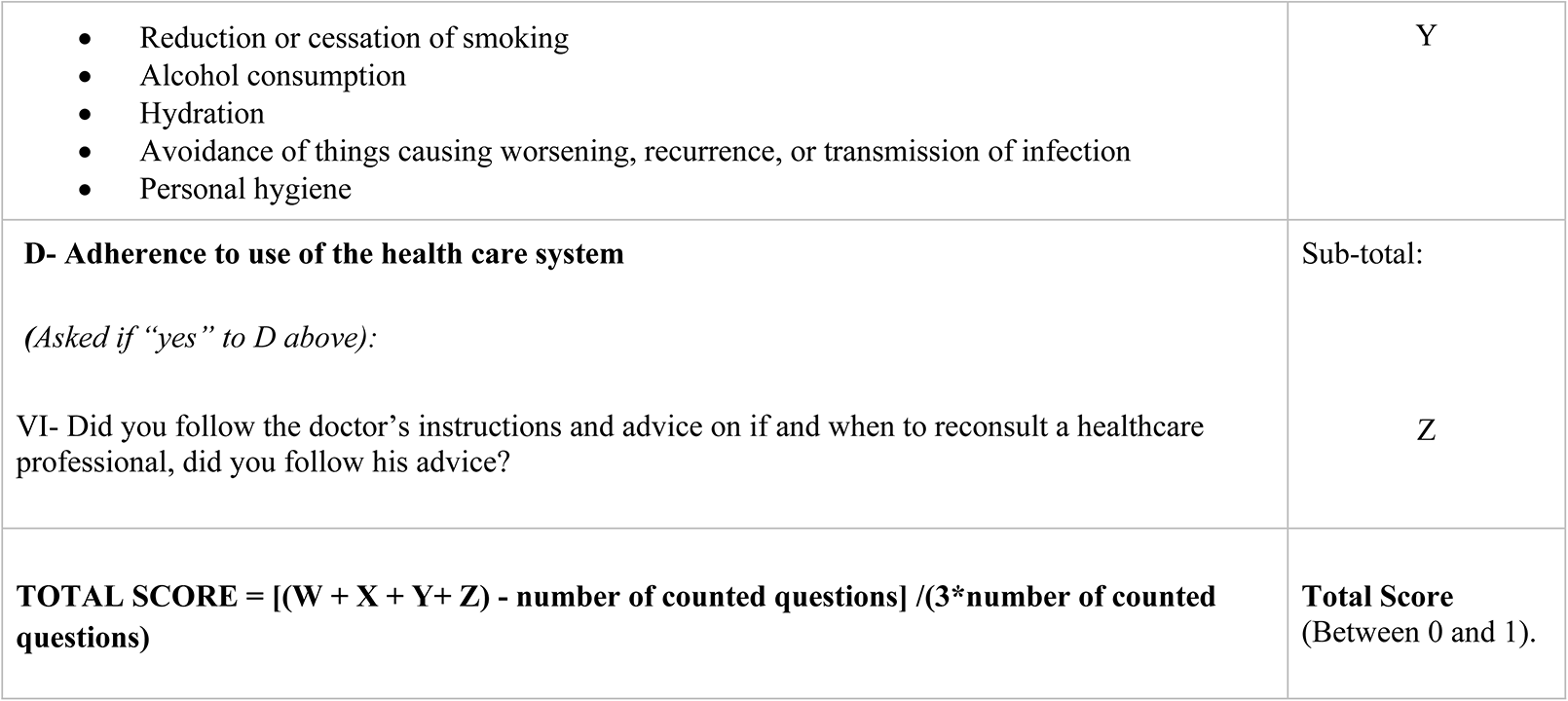
Global Adherence Scale for Acute Conditions (GASAC).

| DOCTOR’S BEHAVIOR |  |  |
| --- | --- | --- |
| Did the doctor give you any of the following: |  |  |
| A- Prescriptions for medications | Yes/No | (Not scored) |
| B- Prescription orders for evaluations, tests or specialized consultation (imagery, laboratory analyses, an appointment with a specialist) | Yes/No |  |
| C- Advice on hygiene, diet, lifestyle, physical activity, regular activities, occupational activity, tobacco or alcohol consumption, hydration, environment | Yes/No |  |
| D- Instructions on when to engage the healthcare system for worsening symptoms | Yes/No |  |
| PATIENT’S BEHAVIOR: THE GASAC QUESTIONNAIRE |  | SCORE |
| Possible answers (Likert scale): No; not at all=1; Partly =2; Mainly=3; Yes, that’s right= 4<br>To each of the following questions: |  |  |
| **A- Adherence to prescription(s) for medication/drugs** (asked if “yes” to A above): |  | Sub-total: |
| I- Have you taken the whole course of prescribed medication? |  | I + II + III = W |
| II- Did you take the prescribed doses? |  |  |
| III- Did you comply with the regimen and conditions (time at which you took the medication, if you were fasted or not, before meals, during meals etc.)? |  |  |
| **B- Adherence to prescription orders for evaluations, tests or specialized consultation** (asked if “yes” to B above) |  | Sub-total: |
| IV- Did you have the additional examinations (imagery, laboratory analyses) or a specialized consultation (an appointment with a specialist) prescribed by the doctor? |  | X |
| **C- Adherence to recommendations and advice** (asked if “yes” to C above): |  | Sub-total: |
| V- Did you follow the advice and recommendations and/or have you changed some habits as a result of the consultation: |  |  |
| Diet and weight control<br>Lifestyle<br>Physical activity |  |  |

|  |  |
| --- | --- |
| • Reduction or cessation of smoking<br>• Alcohol consumption<br>• Hydration<br>• Avoidance of things causing worsening, recurrence, or transmission of infection<br>• Personal hygiene | Y |
| **D- Adherence to use of the health care system**<br>*(Asked if “yes” to D above):*<br>VI- Did you follow the doctor’s instructions and advice on if and when to reconsult a healthcare professional, did you follow his advice? | Sub-total:<br>Z |
| **TOTAL SCORE = [(W + X + Y+ Z) - number of counted questions] /(3\*number of counted questions)** | **Total Score**<br>(Between 0 and 1). |

Table 2 shows the optional supplementary behavioural questions we asked. Supporting information S1 file gives an example of a patient’s replies and how it was scored. Supporting information S2 file contains both the GASAC questionnaire and the supplementary optional questions in French.

**Table 2.**
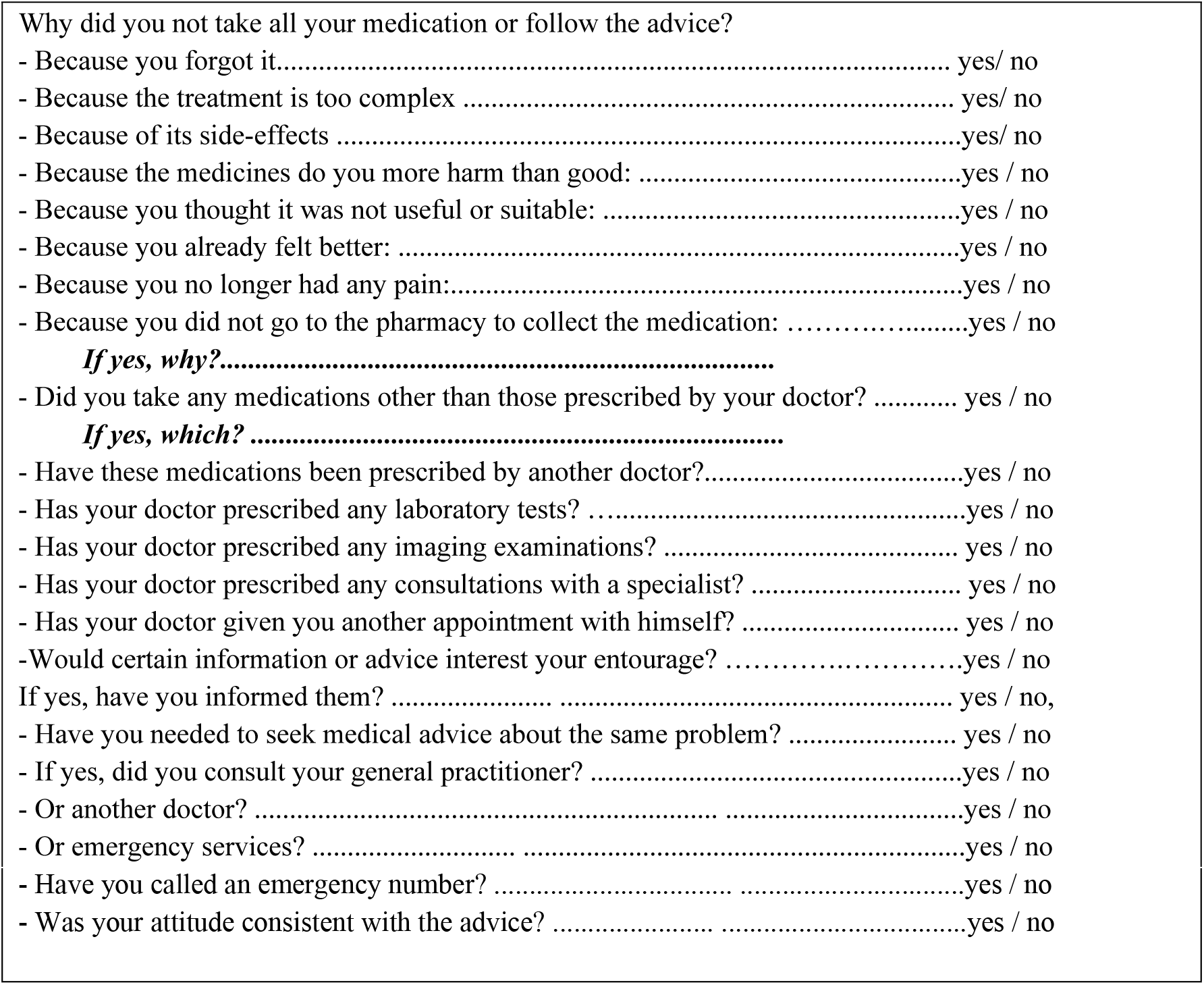
Additional questions about the intentional or non-intentional nature of the patient’s behavior.

### Clinical study

#### Population

The median GASAC score was 0.93 IQR [0.78; 1] (n = 154). Figure 1 shows the patient flow chart.

**Figure 1.**
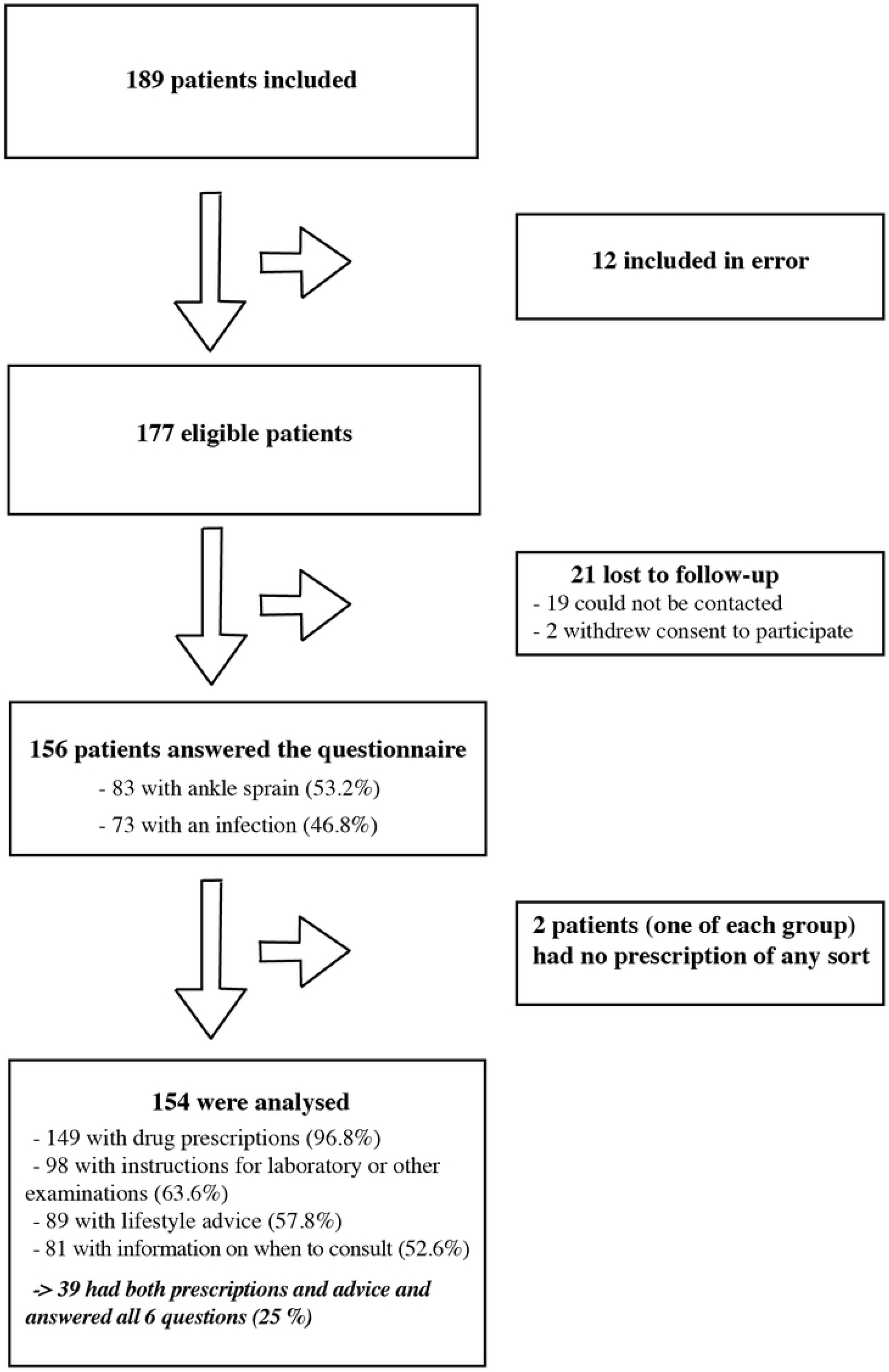
Flow-diagram for the validation of the GASAC scale. Patients lost to follow-up were those who could not be contacted by telephone after 3 attempts.

Table 3 shows the baseline and socio-demographic characteristics of the patients.

**Table 3.** The baseline and socio-demographic characteristics of patients

|  | **All patients**<br>**(n=154)** | **Ankle sprain**<br>**(n=82)** | **Infection (n=72)** |
| --- | --- | --- | --- |
| Female | 93 (60.5%) | 40 (43%) | 53 (57%) |
| Male | 61 (39.6%) | 42 (68.9%) | 19 (31.1%) |
| Age | 36 [23-57] | 27.5 [20-39] | 53.5 [29.5-68.5] |
| Age bands: |  |  |  |
| <40 yrs | 83 (53.9%) | 62 (74.7%) | 21 (25.3%) |
| ≥40 yrs | 71 (46.1%) | 20 (28.2%) | 51 (71.8%) |
| Highest educational level attended: |  |  |  |
| Junior High | 62 (40.3%) | 25 (40.3%) | 37 (59.7%) |
| High school | 38 (24.7%) | 25 (65.8%) | 13 (34.2%) |
| University studies | 54 (35.1%) | 32 (59.3%) | 22 (40.7%) |
| Employed in the health care sector | 22 (14.3%) | 11 (50.0%) | 11 (50.0%) |
| Socio-professional category: |  |  |  |
| 1. Farmers/artisans/tradesmen/<br>Intellectuals/managers | 36 (23.4%) | 22 (61.1%) | 14 (38.9%) |
| 2. Employees/workers | 43 (27.9%) | 26 (60.5%) | 17 (39.5%) |
| 3. Retired | 31 (20.1%) | 5 (16.1%) | 26 (83.9%) |
| 4. Unemployed person | 44 (28.6%) | 29 (65.9%) | 15 (34.1%) |
| Single | 77 (50.0%) | 46 (59.7%) | 31 (40.3%) |
| Living as a couple | 77 (50.0%) | 36 (46.8%) | 41 (53.2%) |

#### Internal validity of the GASAC questionnaire

Cronbach’s alpha coefficient, calculated for the drug subscale was 0.78 (n=149; one-sided Cronbach’s alpha 95% confidence interval 0.72). The other subscales were composed of a single question. It was not relevant to calculate a Cronbach coefficient for global adherence because not all patients were concerned by all the questions.

We calculated the Spearman’s coefficient between the different subscales of the GASAC. There was a correlation between the drug subscale and the use of the health care system (Spearman coefficient = 0.29 with p = 0.01). In contrast, there was no correlation between the drug subscale and the subscale of recommendations and advice, or between the drug subscale and the test and examination subscale.

#### External validity of the GASAC questionnaire

We compared the GASAC and Girerd scores (Figure 2). The Girerd analysis included 149 patients with scores distributed as follows: no patient had a score of less than 3/6; 5 (3.3%) had a score of 3/6; 19 (12.8%) had a score of 4/6; 35 (23.5%) had a score of 5/6 and 90 (60.4%) had a score of 6/6. Using a non-parametric test there was a statistically significant link between the GASAC score and the Girerd score (p<0.01) and between the drug sub-section of the GASAC score and the Girerd score (p<0.01). We also found a statistically significant correlation between the GASAC and satisfaction scores (n= 154; Spearman coefficient=0.22; p<0.01), as well as with the DPC score (n=154; Spearman coefficient = 0.18, p= 0.03).

**Figure 2.**
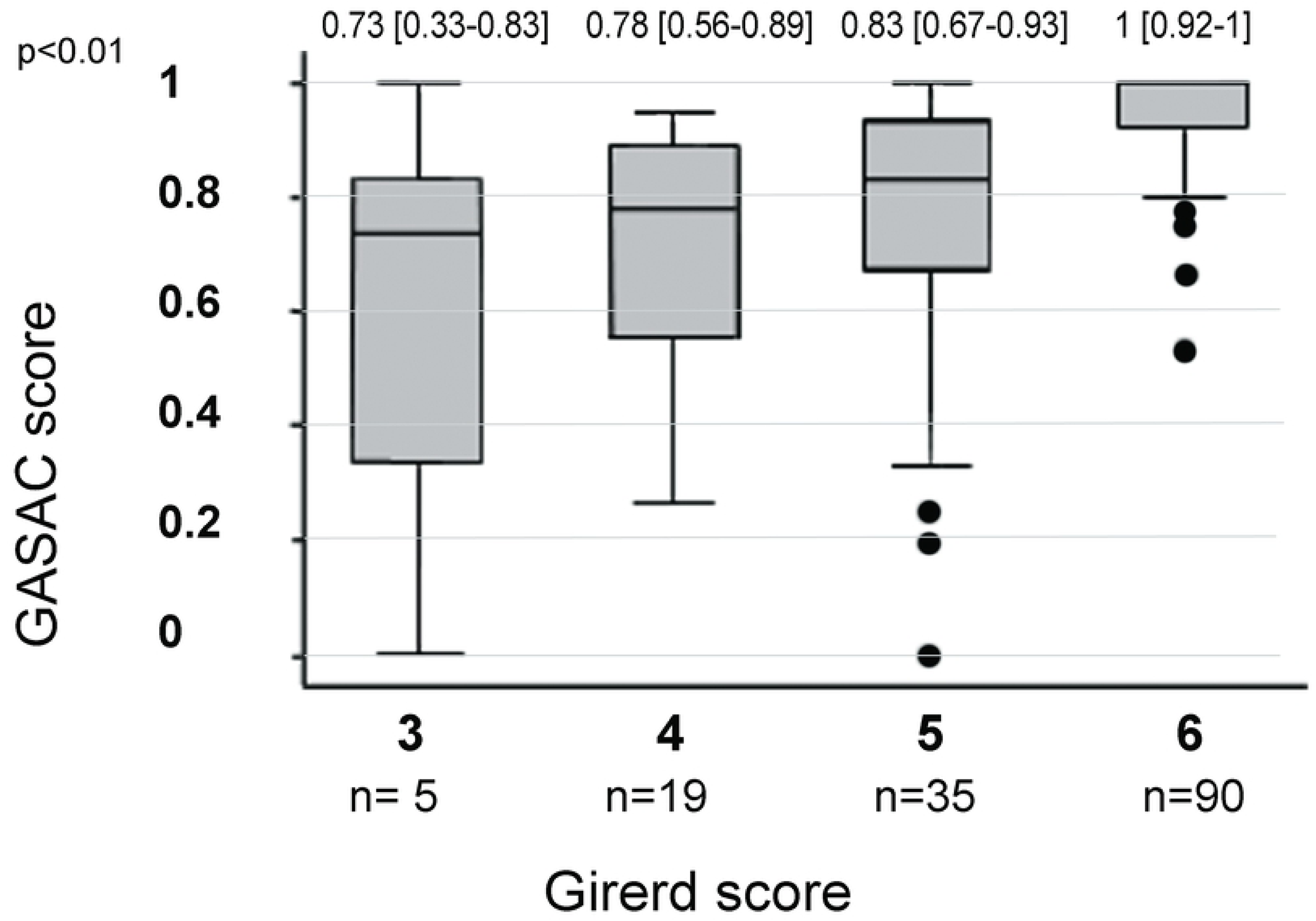
Comparison between GASAC and Girerd adherence scores (n =149). Box plot representing the results by continuous variables using median and IQR [25^th^ and 75 percentile].

#### The determinants of adherence

Using an univariate analysis, we determined that the variables associated (p<0.05) with high adherence were the age-band over 40 years, an infectious pathology (as compared to trauma), high patient satisfaction and a high DPC score.

In the multivariate logistic regression model the explanatory variables for high adherence were the type of acute condition, trauma versus infectious (odds ratio, OR 3.69; IC [1.60-8.52] with p<0.01) and the DPC score (OR 1.06; IC [1.02-1.10] by score point with p<0.01) (Table 4). There was no interaction between these two variables. No individual characteristic, the satisfaction score nor the level of information given, were selected by the manual step-wise selection in the final model. For the multivariate analysis the area under the ROC curve was 0.73. The rate of correctly classified patients was 75.32%, with a Hosmer-Lemeshow chi2 (8) = 7.7 and Prob > chi2 = 0.46.

**Table 4.** Determinants of a good adherence (n=154) after emergency department consultations

| Characteristics of the population studied | High adherence<br>(GASAC Score $\geq 0.8$; $n=115/154$, 74.7%) | OR [95%CI] | p |
| --- | --- | --- | --- |
| **UNIVARIATE ANALYSIS** |  |  |  |
| Age bands<br><40 yrs<br>$\geq 40$ yrs | 54/83 (65.1%)<br>61/71 (85.9%) | 13.28 [1.46-7.34] | < **0.01** |
| Sex<br>Male<br>Female | 41/61 (67.2%)<br>74/93 (79.6%) | 1<br>1.90 [0.91-3.96] | 0.09 |
| Employed in the health care sector<br>No<br>Yes | 99/132 (75.0%)<br>16/22 (72.7%) | 1<br>0.89 [0.32-2.46] | 0.82 |
| Highest educational level attended<br>Junior High<br>High school<br>University studies | 51/62 (82.3%)<br>25/38 (65.8%)<br>39/54 (72.2%) | 1<br>0.41 [0.16-1.06]<br>0.56 [0.23-1.36] | 0.16 |
| Family context<br>Single<br>Living as a couple | 55/77 (71.4%)<br>60/77 (77.9%) | 1<br>1.41 [0.68-2.93] | 0.35 |
| Socio-professional category:<br>Farmers/artisans/tradesmen<br>Intellectuals/managers<br>Employees/Workers/Retirees<br>Unemployed person | 26/36 (72.2%)<br>31/43 (72.1%)<br>28/31 (90.3%)<br>30/44 (68.2%) | 1<br>0.99 [0.37-2.67]<br>3.59 [0.89-14.5]<br>0.82 [0.31-2.17] | 0.15 |
| Type of conditions:<br>Ankle sprain<br>Infection | 53/82 (64.6%)<br>62/72 (86.1%) | 1<br>3.39 [1.51-7.60] | < **0.01** |
| Level of information*<br>0: No information received<br>1: One information received<br>2: The two information received | 24/38 (63.2%)<br>49/64 (76.6%)<br>42/52 (80.8%) | 1<br>1.91 [0.79-4.58]<br>2.45 [0.94-6.36] | 0.15 |
| Satisfaction score (continuous) | GASAC ≥ 0,8: 19 [17-20]<br>GASAC < 0,8: 18 [14-20] | 1.15 by satisfaction<br>score point [1.03-1.28] | **0.01** |
| DPC score (continuous) | GASAC ≥ 0,8: 53,1 [48-56.8]<br>GASAC < 0,8: 50 [37-53] | 1.06 by DPC score<br>point [1.02-1.10] | < **0.01** |
| **MULTIVARIATE ANALYSIS \*\*** |  |  |  |
| DPC score |  | Adjusted OR 1.06<br>[1.02-1.10] by score<br>point | < **0.01** |
| Infection vs trauma |  | Adjusted OR 3.69<br>[1.60-8.52] | < **0.01** |

\* 2 questions were asked about information received: "Has your doctor given you any advice to follow?" And
"Has the doctor given you information on when to reconsult?"
\*\*Final model with manual stepwise logistic regression

### Additional Results

Ninety-eight patients (65.8%) reported having followed the entire treatment; 134 patients (89.9%) reported having taken the prescribed daily doses and 110 (73.8%) said they complied with instructions for the treatment (when and how to take the treatment). Eighty of the 98 patients who had blood tests or radiography prescriptions (81.6%) said they had fully complied with them. Sixty-five patients out of 89 who had been given lifestyle and dietary instructions (73%) said they had followed the advice closely. Sixty-eight (84%) of the 81 patients said they had respected the advice concerning potential reconsultations.

## Discussion

### Characteristics of the GASAC and strengths

The GASAC is a short, patient self-reported questionnaire evaluating four types of patient behaviour following a consultation for an acute condition. The absence of correlations between adherence to drug-prescriptions and adherence to prescription orders for evaluations, tests or specialized consultations, and also with adherence to advice given by the physician, shows that each of the different subscales provides complementary information and are not redundant.

The validity of the intrinsic properties of the drug sub-section of the GASAC score was confirmed by a satisfactory Cronbach’s alpha coefficient^34^ and a high rate of correctly classified patients. For external validity, the study also showed convergence with the Girerd score as well as with well-known determinants of adherence for chronic conditions such as DPC and satisfaction [17,30].

### Clinical results for adherence and comparison with literature

Our median overall score reflects high short-term adherence. This adherence rate is consistent with those reported in the literature for drug adherence for an AC [3,27]. This high rate can be explained by the generally strong motivation of patients seeking ED consultations to follow their treatment, even though the conditions of patient management do not always allow physicians to give detailed explanations. In the literature, the average adherence rates are highly dependent on the context of the study. Nevertheless, adherence rates are higher for acute diseases than for chronic diseases, for which the median adherence is around 40 to 50% [26,39]. In a study using microelectronic monitors to record pill taking Cramer et al. showed that 88% of patients adhere to the treatment prescribed during the first 5 days following the consultation, 86% during the next 5 days, dropping to 67% one month after the consultation [40].

According to a review of reviews on adherence to medical treatment, the median rate of adherence to lifestyle and dietary recommendations is generally lower than for drugs [26]. However, this was not the case in our study: the adherence rate was good for all 4 dimensions of GASAC. This might be explained by the nature of the acute conditions we chose to study.

In univariate analysis, the factors associated with good adherence were age >40 years, an infectious pathology, good satisfaction and quality of DPC. Women tended to be more adherent than men, even if our result wasn’t significant. Patients who lived with a partner also tended to be more adherent. In the literature older and female patients are associated with better adherence [33].

In multivariate analysis, the only determinants of high adherence were infection (versus trauma) and quality of DPC. Whereas the individual characteristics of the patients (age, sex, level of education and professional activity) were not retained in the final model, the DPC score was. This is a major determinant for which measures can be taken to improve, for example through the training of caregivers. The better adherence of patients with an infection might be explained by the risk of deterioration without treatment and the well-known efficacy of antibiotics. Older age and female sex did not appear in the final model because these were also characteristics of patients with infections, as opposed to younger patients presenting with trauma.

### Limitations

It is possible that patients lost to follow-up (who could not be contacted by telephone after 3 attempts) were the least adherent. Our study included patients consulting a hospital emergency department; they may well have had more severe conditions requiring more urgent attention with better adherence rates than others who waited to consult their GP. For minors (n=13, 8%), adherence may well have been improved by parental supervision.

While the emergency physician who included the patient, and the patient were told that the study was aimed to evaluate tools used to assess the quality of care, they weren’t told in detail what was being measured to avoid biases. Moreover, the investigators who made the telephone interviews were completely independent of the emergency departments.

To reduce any social desirability bias (e.g. the patient wanting to be a “good patient”), the interviewers tried to create an atmosphere in which patients did not feel judged, reminding the patient that their answers were anonymous and that the interviewer was not part of the medical team. We considered a telephone questionnaire as being more anonymous and less intimidating for the patient than a face-to-face interview.

It is thought that patients overestimate their level of adhesion when self-reporting by 10 to 20% compared with other methods [21]. Despite this drawback, self-evaluation is frequently used in clinical practice and gives more reproducible results, is more reliable, less expensive and less invasive than direct measurements [28].

To reduce recall bias [20], the telephone interview was made relatively early (7 to 10 days after the consultation) to obtain more reliable answers, however some patients had not completed their treatment or had not been able have all the additional tests required. Although telephone questionnaires have limitations (difficulty understanding questions, call hours and availability vary according to the patient), they allow data to be obtained rapidly, the number of patients lost to follow-up is minimized and they are inexpensive, To optimize accuracy, when measuring adherence it is recommended to combine different types of measures [28,41], direct and indirect [3]. However, this is rarely possible in practice [37,42]. Objective methods would have been further complicated in our study due to the wide range of behaviours we considered (e.g. following a diet, increasing physical activity etc).

For statistical analysis purposes we needed to dichotomize patients into “highly-adherent” or “poorly-adherent” groups, however this led to some simplification and loss of information [21]. For reasons of feasibility, we limited our study to 6 AC corresponding to 2 subgroups of pathology (trauma versus infection). Patients who consulted for an infection outnumbered those seen for a sprained ankle, and posed very different therapeutic issues. However, we cannot generalize our findings to all the pathologies encountered in an ED.

## Conclusion

The Global Adherence Scale for acute conditions is short, patient self-reported, well adapted to the context of an acute condition, independent of the type of condition and takes into account the different aspects of a patient’s behaviour. Such a tool can be useful to assess the quality of healthcare both in clinical research and routine practice. Despite overcrowding in emergency departments, the quality of doctor-patient communication is a major aspect, which can improve adherence. Strategies for improving doctor patient communication should be tested in an interventional study with adherence as the major outcome [23,43].

## Acknowledgments

Pr Girerd for authorization to use his scale.

## Supporting information

**S1 file. An example of a patient’s replies and how it was scored.**

**S2 file. The GASAC questionnaire and the supplementary optional questions in French.**

**S3 file. Protocol in English.**

**S4 file. STROBE Checklist.**

## References

1. Allenet B, Baudrant M, Lehmann A, Gauchet A, Roustit M, Bedouch P, et al. [How can we evaluate medication adherence? What are the methods?]. Ann Pharm Fr 2013;71:135–41.

2. Sabate E. Adherence to long-term therapies. Evidence for action. Geneva: World Health Organization 2003; Available from: whqlibdoc.who.int/publications/2003/9241545992.pdf

3. Ito H. What Should We Do to Improve Patients’Adherence? J Exp Clin Med 2013; 5:127–30.

4. Haynes RB, Mc Donald HP, Garg AX. Helping patients follow prescribed treatment: clinical applications. J Am Med Assoc 2002; 288: 2880–3.

5. Gauchet A, Tarquinio C, Fischer G. Psychosocial predictors of medication adherence among persons living with HIV. Int J Behav Med 2007;14:141–50.

6. El-Saifi N, Moyle W, Jones C, Tuffaha H. Medication Adherence in older patients with dementia: a systematic review. J Pharm Pract. 2018 Jun;31(3):322–334.

7. Virjens B, De Geest S, Hughes DA, Przemyslaw K, Demonceau J, Ruppar T, et al. A new taxonomy for describing and defining adherence to medications: new taxonomy for adherence to medications. Br J Clin Pharmacol 2012; 3: 691–705.

8. McDonald HP, Garg AX, Haynes RB. Interventions to Enhance Patient Adherence to Medication Prescriptions: Scientific Review. J Am Med Assoc 2002; 288: 2868–2879.

9. Raynor DK, Blenkinsopp A, Knapp P, Grime J, Nicolson DJ, Pollock K, et al. A systematic review of quantitative and qualitative research on the role and effectiveness of written information available to patients about individual medicines. Health Technol Assess Winch Engl 2007;11: 1–160.

10. Girerd X, Hanon O, Anagnostopoulos K, Ciupek C. Assessment of antihypertensive compliance using a self-administered questionnaire: development and use in a hypertension clinic. Presse Med. 2001 Jun 16-23;30(21):1044–8.

11. Girerd X, Radauceanu A, Achard JM. Evaluation of patient compliance among hypertensive patients treated by specialists. Arch Mal Cœur Vaiss. 2001 Aug;94(8):839–42.

12. Yeowell G, Smith P, Nazir J, Hakimi Z, Siddiqui E, Fatoye F. Real-world persistence and adherence to oral antimuscarinics and mirabegron in patients with overactive bladder (OAB): a systematic literature review.BMJ Open. 2018 Nov 21;8(11):e021889.

13. Murage MJ, Tongbram V, Feldman SR, Malatestinic WN, Larmore CJ, Muram TM, et al. Medication adherence and persistence in patients with rheumatoid arthritis, psoriasis, and psoriatic arthritis: a systematic literature review. Patient Prefer Adherence. 2018 Aug 21;12:1483–1503.

14. Haynes RB, Taylor WD, Sackett DL. Compliance in health care. Baltimore: Johns Hopkins University Press 1979; 1–15.

15. Sustersic M, Gauchet A, Foote A, Bosson JL. How best to use and evaluate Patient Information Leaflets given during a consultation: a systematic review of literature reviews. Health Expect 2016; 1–12.

16. Sustersic M, Tyrant J, Tissot M, Gauchet A, Foote A, Vermorel C, et al. Impact of patient information leaflets on doctor–patient communication in the context of acute conditions: a prospective controlled before–after study in two French emergency departments. BMJ Open 2019;9:e024184.

17. Vermeire E, Hearnshaw H, Van Royen P, Denekens J. Patient adherence to treatment: three decades of research. A comprehensive review. J Clin Pharm Ther 2001;26:331–42.

18. Lavsa SM, Holzworth A, Ansani NT. Selection of a validated scale for measuring medication adherence. J Am Pharm Assoc 2011; 51:90–4.

19. Lehmann A, Aslani P, Ahmed R, Celio J, Gauchet A, Bedouch P, et al. Assessing medication adherence: options to consider Int J Clin Pharm 2013; 36: 55–69.

20. Garfield S, Clifford S, Eliasson L, Barber N, Willson. Suitability of measures of selfreported medication adherence for routine clinical use: A systematic review. BMC Med Res Methodol 2011;11:149.

21. Voils CI, Hoyle RH, Thorpe CT. Improving the measurement of self-reported medication nonadherence. J Clin Epidemiol 2011;64:250–4.

22. French society of general practitioners (SFMG). http://www.sfmg.org/theorie_pratique/outils_de_la_demarche_medicale/le_dictionnaire_des_resultats_de_consultation-drc/

23. Ackermann S, Bingisser MB, Heierle A. Discharge communication in the emergency department: physicians underestimate the time needed. Swiss Med Wkly 2012;142: 1–6.

24. Kripalani S, Jackson AT, Schnipper JL, Coleman EA. Promoting effective transitions of care at hospital discharge: a review of key issues for hospitalists. J Hosp Med. 2007;2(5):314–23. Epub 2007/10/16

25. Laufs U, Rettig-Ewen V, Böhm M. Strategies to improve drug adherence. Eur Heart J 20111; 32:264–8.

26. Dulmen SV, Sluijs E, Dijk L van, Ridder D de, Heerdink R, Bensing J. Patient adherence to medical treatment: a review of reviews. BMC Health Serv Res 2007;7:55.

27. Nieuwlaat R, Wilczynski N, Navarro T, Hobson N, Jeffery R, Keepanasseril A, et al. Interventions for enhancing medication adherence. Cochrane Database Syst Rev 2014: CD000011.

28. DiMatteo MR. Variations in patients’ adherence to medical recommendations: a quantitative review of 50 years of research. Med Care 2004;42:200–9.

29. Nicolson D, Knapp P, Raynor D, Spoor P, Knapp P. Written information about individual medicines for consumers. Cochrane Database Syst Rev Online 2009;(2):CsD002104.

30. Simmons S, Sharp B, Fowler J, Fowkes H, Paz-Arabo P, Dilt-Skaggs MK, et al. Mind the (knowledge) gap: The effect of a communication instrument on emergency department patients’ comprehension of and satisfaction with care. Patient Educ Couns 2015; 98: 257–262.

31. Bos N, Sizmur S, Graham C, Van Stel HF. The accident and emergency department questionnaire: a measure for patients’ experiences in the accident and emergency department. BMJ Qual Saf 2013;22:139–146.

32. Tarquinio C, Fischer G, Grégoire A. Compliance in HIV-positive patients: validation of a French scale and measurement of psychosocial variables. Rev Int Psychol Soc 2000;13(2):61–91.

33. Morisky DE, Ang A, Krousel-Wood M, Ward HJ. Predictive validity of a medication adherence measure in an outpatient setting. J Clin Hypertens. 2008 May;10(5):348–54.

34. Ruiz MA, Pardo A, Rejas J, Soto J, Villasante F, Aranguren JL. Development and validation of the “Treatment Satisfaction With Medicines Questionnaire” (SATMED-Q). Value Health. Oct 2008; 11(5): 913–926.

35. AHRQ. Care Coordination Measures Atlas [Internet]. Agency for Healthcare Research and Quality; 2010. Available from: http://www.cshcndata.org/docs/drc/ahrq-care-coordination-atlas-dec-2010.pdf

36. Richard C, Lussier MT. [Professional communication in Health]. 2nd edition.Saint-Laurent, Quebec, Canada: Editions du Renouveau pedagogique; 2016.

37. Garratt AM, Bjærtnes A, Krogstad U, Gulbrandsen P. The OutPatient Experiences Questionnaire (OPEQ): data quality, reliability, and validity in patients attending 52 Norwegian hospitals. Qual Saf Health Care 2005;14:433–437.

38. Bland JM, Altman DG. Statistics notes: Cronbach’s alpha. BMJ 1997; 314:572.

39. Baudrant M. [Reflections on the role of the pharmacist in patient therapeutic education]. J Pharm Clin 2008; 27:201–4.

40. Cramer JA, Scheyer RD, Mattson RH. Compliance declines between clinic visits. Arch Intern Med 1990;150:509e10.

41. Osterberg L, Blaschke T. Adherence to medication. N Engl J Med 2005; 353:487–97.

42. Fossli Jensen B, Dahl FA, Safran DG, Garratt AM, Krupat E, Finset A et al. The ability of a behaviour-specific patient questionnaire to identify poorly performing doctors. BMJ Qual Saf 2011; 20: 885–893.

43. Hansoti B, Aluisio AR, Barry MA, Davey K, Lentz BA, Modi P, et al. Global Health and Emergency Care: Defining Clinical Research Priorities. Acad Emerg Med 2017; 24(6): 742–753.

